# An engineered serum albumin-binding AAV9 capsid achieves improved liver transduction after intravenous delivery in mice

**DOI:** 10.1101/694513

**Authors:** Quan Jin, Chunping Qiao, Jianbin Li, Juan Li, Xiao Xiao

## Abstract

Recombinant adeno-associated viral (AAV) vectors are frequently used to deliver nucleic acids for in vivo applications and are currently the leading platform for therapeutic gene delivery in gene therapy clinical trials. Presently, there is a need for improved AAV vectors with optimized transduction efficiency in target tissues. In these studies, an engineered albumin-binding consensus domain (ABDCon) was incorporated into the AAV9 capsid via fusion to the N-terminus of the AAV9 VP2 capsid protein to generate a variant AAV9 capsid with albumin-binding properties. The variant capsid, called AAV9-ABDCon, formed viable genome-containing vector particles and exhibited binding to human serum albumin. The AAV9 capsid, on the other hand, was not found to bind to human serum albumin by the methods used in this study. Following intravenous administration, the modified AAV9-ABDCon vector was found to achieve higher levels of transduction in liver tissue compared to AAV9. These findings suggest that serum albumin-binding may be a potential method to augment AAV-mediated liver-directed gene delivery.

## INTRODUCTION

Recombinant AAV vectors are frequently used for in vivo gene transfer applications and are currently the most commonly used vector for therapeutic gene delivery in gene therapy clinical trials (1). The liver is affected in many genetic diseases, and considerable research effort has been directed toward leveraging AAV vectors to deliver therapeutic transgenes to target genetic diseases afflicting the liver (2–5).

However, the clinical development of AAV vectors for liver-directed gene delivery has been hindered by several obstacles. First, existing naturally-derived AAV serotypes may not achieve anticipated transduction efficiency in human hepatocytes (6). Another issue, as noted in earlier clinical trials, is clearance of transduced hepatocytes by a capsid-specific cytotoxic T-lymphocytes (CTLs) and subsequent loss of transgene expression (7,8). This capsid-specific CTL response may be dose-limiting as the CTL response is amplified at higher vector loads (9). One way to address these issues is to identify or engineer novel AAV capsids with improved transduction efficiency in the liver, which would not only mediate more efficient gene delivery in vivo but also allow for a reduction in the vector dose required to achieve therapeutic effect, thus lowering the overall capsid burden.

Previous studies have reported that pre-incubation of AAV8 with human serum albumin (HSA) increases AAV transduction in vitro and in vivo, and this effect was also observed across numerous AAV serotypes (10,11). However, we were not able to detect binding of AAV9 to HSA. Thus, to ascertain the impact of AAV albumin-binding on vector tropism and transduction, we designed a modified AAV9 capsid capable of binding to serum albumin in systemic circulation. This was achieved via insertion of a non-natural albumin binding domain consensus sequence (ABDCon) peptide derived from the *Streptococcus* spp. G148 protein G into the AAV9 capsid. The ABDCon peptide is a three-helix bundle 45 amino acids in length, and has been shown to bind to serum albumin from several species, including human and mouse, with high affinity (12).

The AAV2 VP2 N-terminal region has been well-characterized as a site that is amenable to large peptide insertions (13,14). In this study, we construct a modified serum albumin-binding AAV9 capsid via insertion of the engineered ABDCon peptide at the N-terminus of the AAV9 VP2 capsid protein (AAV9-ABDCon). In addition, evaluation of in vivo tropism following systemic vector administration revealed that AAV9-ABDCon achieves higher levels of transgene expression in the liver when compared to unmodified AAV9.

## MATERIALS & METHODS

### Generation of modified AAV9-ABDCon rep/cap plasmids

The AAV9 rep/cap plasmid was modified to incorporate the ABDCon peptide insertion using methods that have been previously described (Supplementary Methods) (13). Purified vectors were titered by quantitative DNA dot blot, and capsids were analyzed by western blot and denaturing alkaline agarose gel electrophoresis (Supplementary Methods).

### Capsid binding ELISA

AAV vector binding to serum albumin was evaluated by enzyme-linked immunosorbent assay (ELISA). Wells on a MICROLON® medium-binding ELISA plate (Greiner Bio-One; Frickenhausen, Germany) were coated with 2 mg/ml human serum albumin (HSA) for 1 hour at room temperature. Wells were blocked with 20 mg/ml BSA in PBS. 2-fold serial dilutions of 1×10^10^ vg of purified AAV vectors were added to the HSA-coated wells for 1 hour and unbound vectors were washed away with 0.1% PBS-Tween 20. The primary antibody used was mouse monoclonal anti-AAV9 intact particle (1:100; ADK9; American Research Products; Waltham, MA) and the secondary antibody used was HRP-conjugated goat polyclonal anti-mouse IgA (1:500; A4789; Sigma Aldrich; St. Louis, MO). HRP was detected with 1-Step Ultra TMB-ELISA Substrate Solution (Thermo Scientific; Rockford, IL) and the reaction was stopped with 2N sulfuric acid. Absorbance was measured at 450 nm.

### Immuno-transmission electron microscopy (TEM)

AAV vector samples were adsorbed onto glow-discharged formvar/carbon-coated nickel grids (FCF400-Ni-SC; Electron Microscopy Sciences; Hatfield, PA) for 2 min. Grids were blocked in blocking solution (30 mM Tris-HCl, 150 mM NaCl, 0.1% BSA, 1% gelatin, pH 8.0) prior to incubation with HSA (1 mg/ml in blocking solution) for 30 min. Grids were washed, incubated in primary followed by secondary antibody for 30 min each, and stained in 2% uranyl acetate for 30 s. Images were taken on a JEOL 1230 TEM (Peabody, MA) at 80,000X to 150,000X magnification. The antibodies used were mouse anti-human HSA (1:500; clone HSA-11, A6684; Sigma Aldrich; St. Louis, MO) and 10 nm colloidal gold-conjugated goat anti-mouse IgG (1:40; G7652; Sigma Aldrich; St. Louis, MO). These methods were partially adapted from a protocol by Rojas and colleagues (15).

### Mice and vector administration

For gene delivery of LacZnls, 6-week-old male and female C57BL/6J mice were administered AAV9-CB-LacZnls (n=5) or AAV9-ABDCon-CB-LacZnls (n=5) by intravenous tail vein injection at a dose of 1×10^13^ vg/kg. For follistatin gene delivery, 6-week-old male C57BL/6J mice were administered vehicle (n=3), AAV9-TBG-Fstn (n=4) or AAV9-ABDCon-TBG-Fstn (n=4) by tail vein injection at a dose of 5×10^12^ vg/kg. At the end of the trials, mice were anesthesized with 2.5% 2,2,2-tribromoethanol (Avertin) at a dose of 350 mg/kg and sacrificed. Tissues were excised, flash-frozen in liquid nitrogen-cooled 2-methylbutane and stored at −80°C.

### LacZ enzymatic activity

For X-gal staining of tissue sections, 10 μm cryosections were prepared. Cryosections were fixed in X-gal fixation buffer (2% formaldehyde, 0.2% glutaraldehyde in PBS, pH 7.3) and stained in X-gal staining buffer (1 mg/ml X-gal, 5 mM K_3_Fe(CN)_6_, 5 mM K_4_Fe(CN)_6_-3H_2_O, mM MgCl_2_ in PBS, pH 7.3). Tissues were incubated in staining buffer overnight except for liver sections, which were incubated for <20 min. β-galactosidase enzymatic activity in tissue lysates was measured using the Galacto-Star System (Applied BioSystems; Bedford, MA) according to the manufacturer’s instructions. RLU readings were normalized to total protein content as measured by Pierce BCA Protein Assay Kit (ThermoFisher Scientific; Waltham, MA). Vector genome copies were measured by qPCR (Supplementary Methods).

### Statistical analysis

Values are presented as mean ± SEM. Student’s t test was used for comparison between two groups. One-way ANOVA was used for comparison of three or more groups. All statistical analyses were conducted in GraphPad Prism (GraphPad Software; La Jolla, CA). p<0.05 was considered statistically significant.

## RESULTS

### Construction of AAV9-ABDCon

For surface display of the ~5.4 kDa ABDCon peptide, The N-terminus of AAV9 VP2 was selected as the peptide insertion site (13). The relatively lower abundance of the AAV9 VP2 compared to the AAV9 VP3 would theoretically allow for sufficient display of the inserted ABDCon peptide without superfluous peptide display, which may lead to undesirably excessive serum albumin binding and possible masking of receptor binding epitopes on the capsid surface. In order to display the ABDCon peptide on VP2 specifically, two separate AAV cap gene variants had to be generated. The first variant, called pVP2-ABDCon was designed to supply the VP2-ABDCon fusion protein gene. A second cap gene variant was generated to produce the unmodified VP1 and VP3 capsid proteins (pVP1,3). Altogether, these mutations permit translation of unmodified AAV9 VP1 and VP3 and the VP2-ABDCon fusion protein to form the AAV9-ABDCon vector capsid (Figure 1).

**Figure 1.**
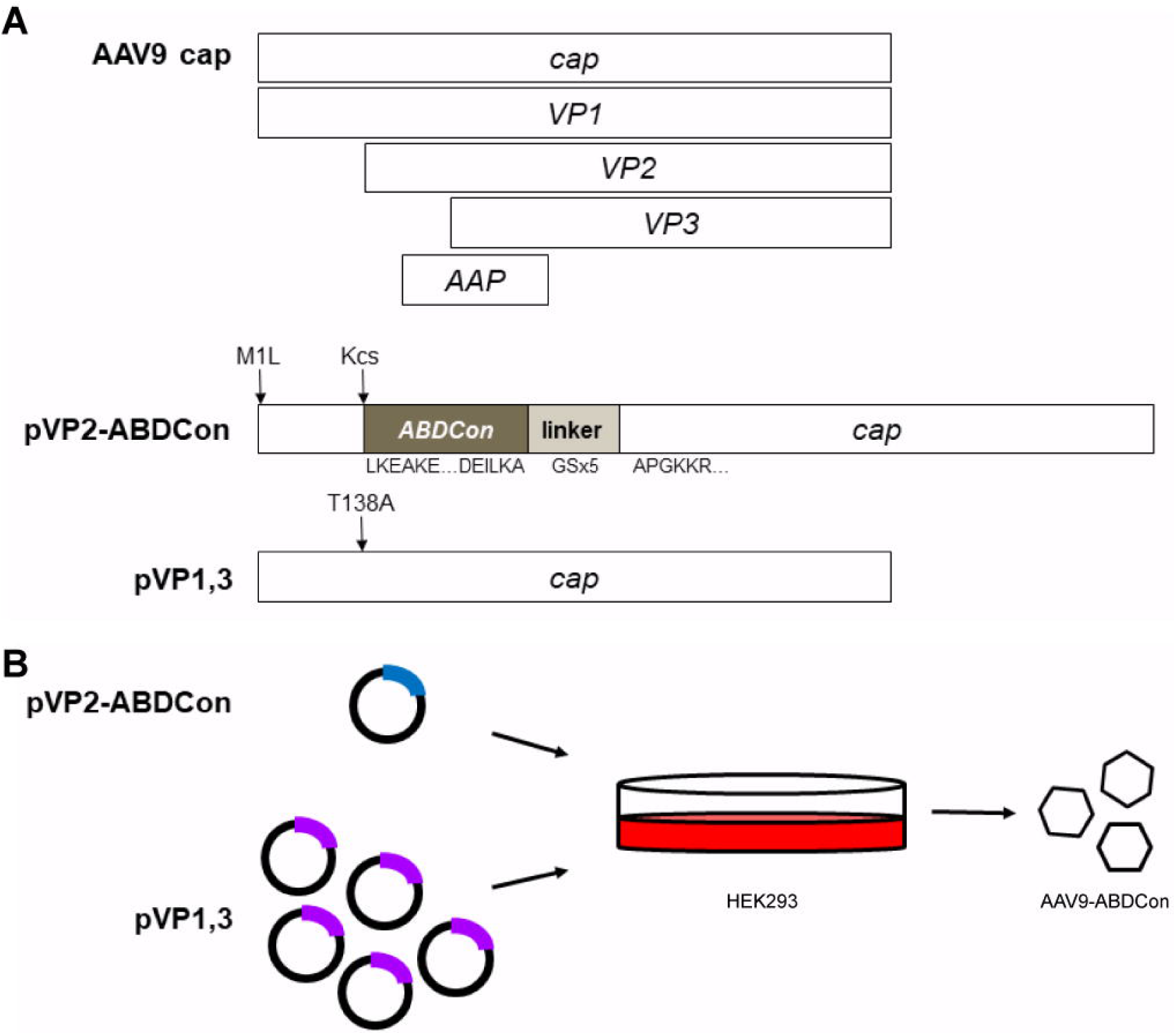
AAV9-ABDCon vector construction. **(A)** Depiction of the AAV9 cap gene with the location of the VP1, VP2 and VP3 coding sequences (top). To generate the modified AAV9-ABDCon capsid, the ABDCon peptide sequence preceded by a Kozak consensus sequence (Kcs) is incorporated at the N-terminus of the AAV9 VP2 with a flexible GSx5 linker (pVP2-ABDCon). To prevent undesired expression of a VP1-ABDCon fusion protein, the VP1 translation start site is mutated (M1L). pVP2-ABDCon supplies the VP2-ABDCon fusion protein. Unmodified VP1 and VP3 are supplied by a second cap sequence with the VP2 translation start site removed (T138A; pVP1,3). **(B)** AAV9-ABDCon capsids are produced by plasmid transfection of pVP1,3 (purple) and pVP2-ABDCon (blue) in a 5:1 molar ratio in adherent HEK293 cells.

### AAV9-ABDCon vector characterization

To generate AAV9-ABDCon vector capsids, a modified version of the triple plasmid transfection method was used where the single rep/cap plasmid is replaced with the two plasmids encoding for the variant VP1,3, supplying VP1 and VP3, and VP2-ABDCon, supplying the modified ABDCon-containing VP2, cap genes in a 5:1 ratio (16,17). To assess whether insertion of the ABDCon peptide at the N-terminus of AAV9 VP2 results in the formation of viable vector particles, AAV9 and AAV9-ABDCon vectors were purified from HEK293 cells 48 h post-transfection and vector titers were determined by quantitative DNA dot blot. The titers for AAV9-ABDCon were approximately 70.7% of unmodified AAV9 titers on average (p=0.1875; Figure 2A).

**Figure 2.**
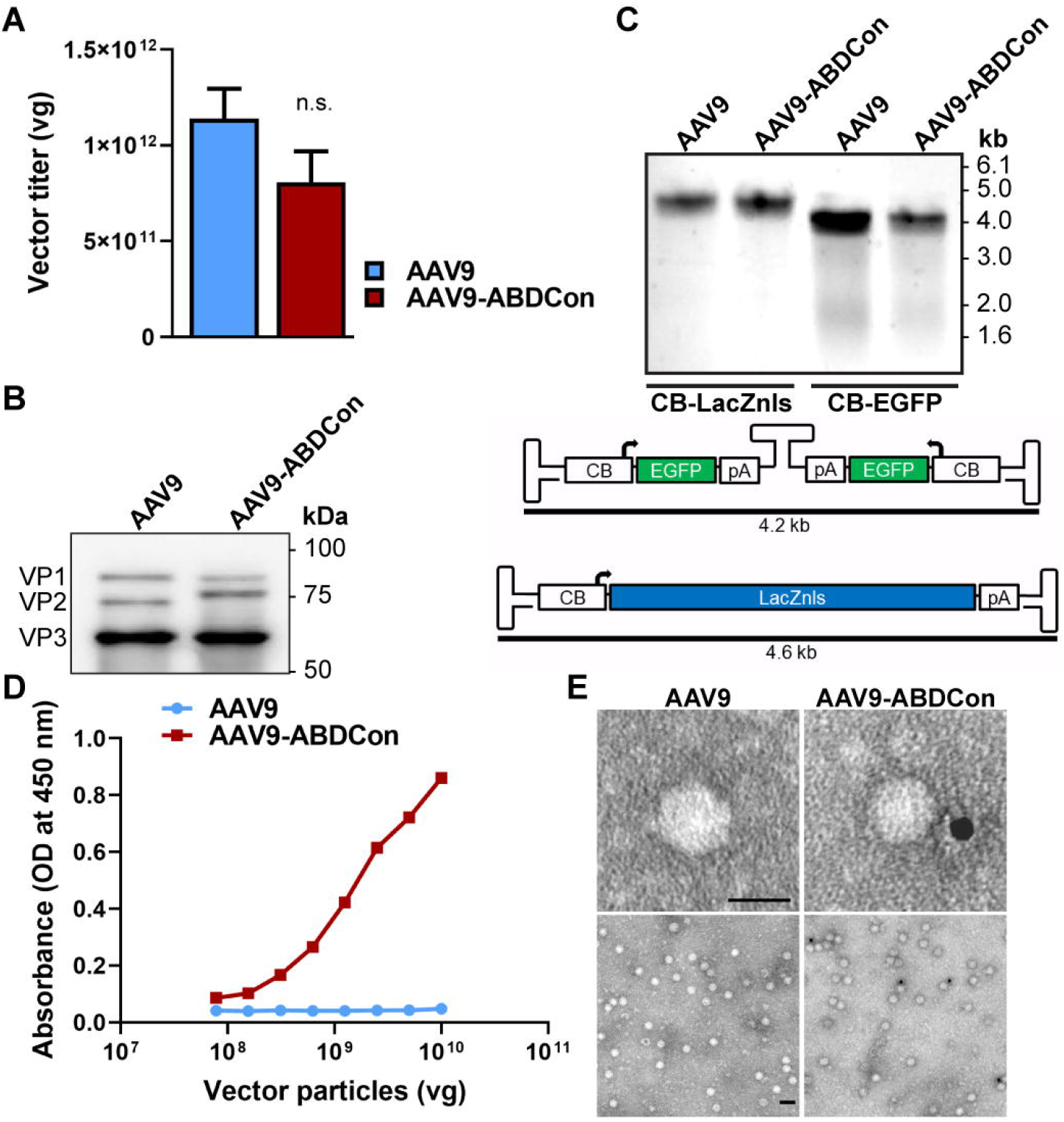
AAV9-ABDCon vector characterization. **(A)** Average quantitative DNA dot blot titers of multiple AAV9 (n=4) and AAV9-ABDCon (n=4) vector preparations. Data represents titer from a single 15-cm plate transfection. **(B)** Western blot of denatured AAV9 and AAV9-ABDCon capsids. VP1, VP2 and VP3 are visible at the expected sizes. The VP2-ABDCon fusion protein migrates slightly higher (79 kDa) than the unmodified AAV9 VP2 (72 kDa). An anti-AAV VP1/VP2/VP3 mouse monoclonal B1 antibody was used to detect AAV VPs. **(C)** Denaturing alkaline agarose gel electrophoresis of AAV9- and AAV9-ABDCon-packaged vector genomes. Both ssAAV and scAAV preparations are depicted. **(D)** ABDCon peptide surface display was assessed by capsid-binding ELISA in HSA-coated ELISA wells. AAV9-ABDCon capsids display binding to HSA whereas unmodified AAV9 exhibit no detectable binding. **(E)** Representative immuno-TEM images of AAV9 and AAV9-ABDCon vectors incubated in HSA. HSA was detected with a mouse anti-HSA primary antibody and a 10 nm colloidal gold-conjugated anti-mouse IgG secondary antibody. Scale bar represents 25 nm (top) and 50 nm (bottom). Error bars represent mean◻±◻SEM. n.s. not significant

Incorporation of the ABDCon peptide onto the AAV9 VP2 N-terminus was confirmed by western blot. As expected, the bands corresponding to VP1 (87 kDa) and VP3 (61 kDa) were at the same position in both denatured AAV9 and AAV9-ABDCon capsid samples. However, the band corresponding to VP2 in AAV9-ABDCon capsids (79 kDa) was shifted further up on the blot in comparison to the VP2 band of AAV9 capsids (73 kDa; Figure 2B). Vector genome integrity was assessed by alkaline agarose gel electrophoresis, and packaging of either a 4.6 kb single-stranded AAV (ssAAV; pAAV2.1-CB-LacZnls) expression cassette or a 2.1 kb self-complementary AAV (scAAV; pEMBOL-ds-CB-EGFP) expression cassette was evaluated. The presence of a distinct band at 4.6 kb and 4.2 kb corresponding to the full-length ssAAV and scAAV genomes, respectively, indicates that both ssAAV and scAAV expression cassettes are packaged into the AAV9-ABDCon capsids (Figure 2C).

AAV9-ABDCon binding to HSA was determined by capsid-binding ELISA. Purified vectors were applied to wells on an ELISA plated coated with human serum albumin. A primary antibody (ADK9) that specifically recognizes intact AAV9 capsids, but not unincorporated AAV capsid VPs, was used to identify samples that were bound to HSA. A positive signal was observed in wells containing purified AAV9-ABDCon samples, which demonstrates that the AAV9-ABDCon capsid binds HSA. On the contrary, samples of unmodified AAV9 capsid did not elicit a signal greater than background, suggesting that unmodified AAV9 capsid does not bind human serum albumin with high affinity. In addition, free VP2-ABDCon subunit also did not elicit a signal, indicating that the ADK9 antibody recognizes intact AAV9 capsids specifically (Figure 2D). HSA-binding AAV9-ABDCon particles were detectable with immuno-TEM (Figure 2E).

### Evaluation of AAV9-ABDCon tissue transduction following intravenous administration

To assess the impact of ABDCon peptide insertion onto the AAV9 VP2 N-terminus on in vivo transduction and tropism, male and female 6-week old C57BL/6J mice were injected intravenously with 1×10^13^ vg/kg of AAV9 or AAV9-ABDCon vectors carrying a nuclear-localizing β-galactosidase (β-gal; LacZnls) reporter transgene under the transcriptional regulation of a chicken β-actin (CB) promoter. LacZnls protein expression across various tissues was assessed 4 weeks post-injection. X-gal staining was more intense in the livers of male mice treated with the AAV9-ABDCon vector. As reported previously, relatively little transgene expression was observed in the livers of female mice across all groups (Figure 3A). A higher level of β-gal enzymatic activity was detected in liver tissue from mice administered the AAV9-ABDCon vector (+319%; p=0.0008; Figure 3B). This difference was associated with a higher vector genome copy number (VGCN) per diploid genome in the liver (20.7 ± 4.60 and 34.8 ± 3.84 for AAV9 and AAV9-ABDCon, respectively; p=0.0464; Figure 3C). β-gal expression in the heart, pancreas, skeletal muscle and spleen were generally unchanged, and there was no statistically significant difference seen in β-gal enzymatic activity or vector copies in the heart, pancreas or skeletal muscle between the groups (Figure 3D-F). Unlike what was observed in the liver, there was no sex-specific differences in β-gal protein expression in the heart, pancreas or skeletal muscle (data not shown). Increased liver transduction was not observed with non-albumin-binding AAV9 vectors containing a VP2 N-terminal peptide insertion, indicating that the increase in liver transduction was not due to random peptide insertion at the VP2 N-terminus (data not shown). Altogether, these results suggest that AAV9-ABDCon achieves increased liver transduction over AAV9, but transduction in peripheral tissues is unchanged.

**Figure 3.**
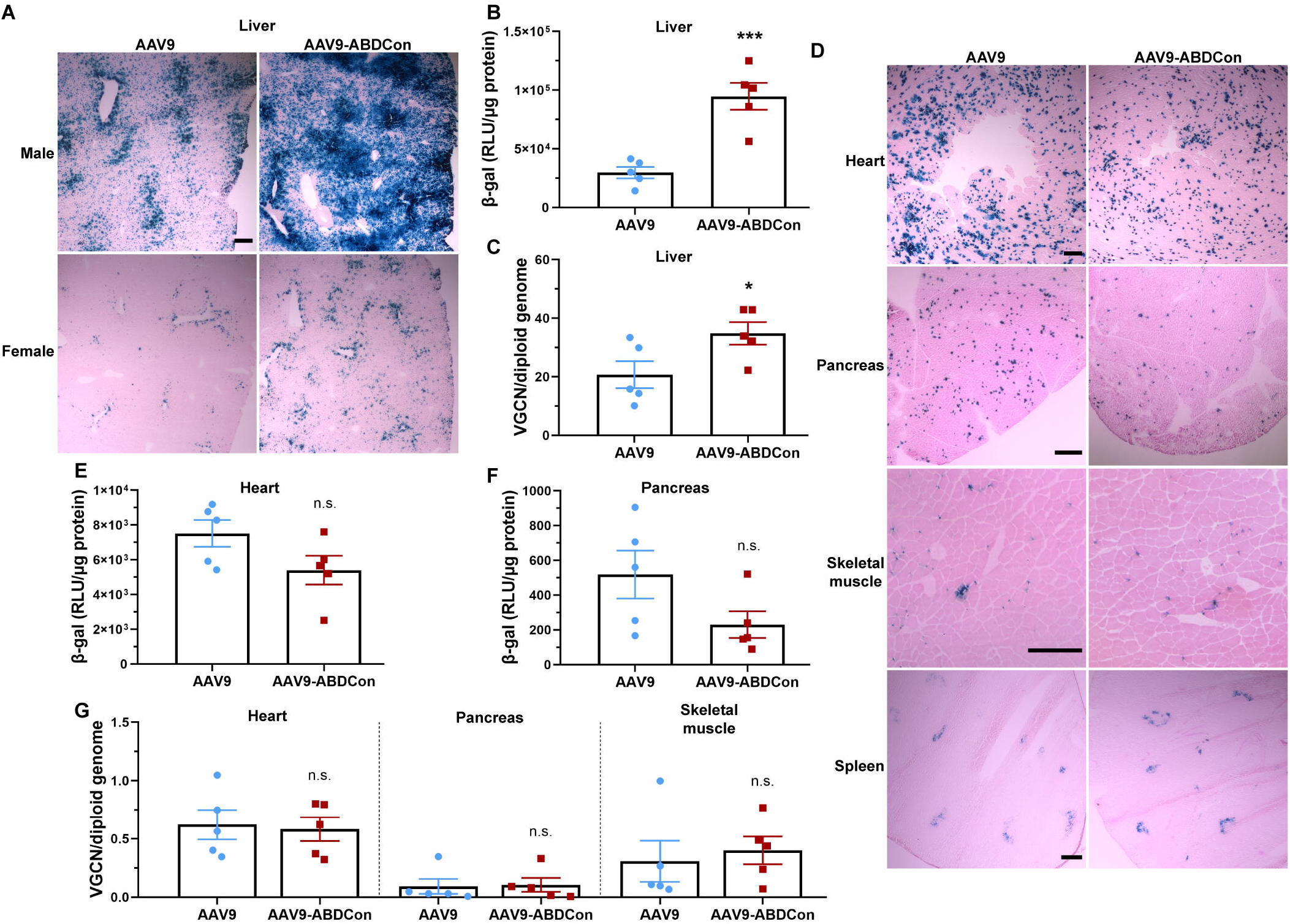
Quantification of AAV9-ABDCon tissue transduction efficiency in mice. Male and female 6-week-old C57BL/6J mice were treated with AAV9-CB-LacZnls (n=5) or AAV9-ABDCon-CB-LacZnls (n=5) by tail vein injection at a dose of 1×10^13^ vg/kg. **(A)** Representative X-gal stained (20 min incubation) liver sections from male and female mice. Staining was more intense in liver sections from male mice. Scale bars represent 200 μm. **(B)** β-gal enzymatic activity quantification in liver tissue homogenates from treated male mice. **(C)** AAV vector genome content in liver tissue. **(D)** Representative X-gal stained (overnight incubation) heart, pancreas, skeletal muscle (gastrocnemius) and spleen sections. **(E)** β-gal enzymatic activity quantification in heart and **(F)** pancreas. Scale bars represent 200 μm. **(G)** AAV vector genome content in heart, pancreas and skeletal muscle (gastrocnemius). All error bars represent mean◻±◻SEM. *p◻<◻0.05; ***p◻<◻0.001; n.s. not significant, VGCN: vector genome copy number

Next, we aimed to evaluate whether the observed increase in liver transduction by AAV9-ABDCon would translate to a more potent transgene-mediated effect following liver-directed gene delivery of follistatin (Fstn). Fstn overexpression has been shown to induce skeletal muscle hypertrophy via inhibition of transforming growth factor β (TGFβ) ligands such as myostatin and the activins (18,19). To ensure that the effect would be limited primarily to Fstn expression and secretion from the liver, we utilized the liver-specific thyroxine-binding globulin (TBG) promoter to drive transgene expression (Figure 4A) (20). 6-week-old male C57BL/6J were injected intravenously with AAV9-TBG-Fstn or AAV9-ABDCon-TBG-Fstn at a dose of 5×10^12^ vg/kg or vehicle. As expected, the average bodyweights of mice treated with both AAV9-TBG-Fstn and AAV9-ABDCon-TBG-Fstn exceeded vehicle-treated controls starting from 2 weeks post-injection (+17.6% and +20.4%; p=0.0053 and p=0.0023 for AAV9 and AAV9-ABDCon, respectively, relative to vehicle; Figure 4B). This increase in bodyweight was associated with skeletal muscle hypertrophy in vector-treated mice, with the largest increase in hindlimb muscle weight measured in mice treated with the AAV9-ABDCon vector (+19.7%, +17.5% and +16.2%; p=0.0036, p=0.0014 and p=0.0020 for tibialis anterior, gastrocnemius and quadriceps muscle weight, respectively, relative to AAV9; Figure 4C-D). Western blot of liver tissue homogenates revealed increased Fstn protein expression in vector-treated mice, with a significantly higher level of Fstn protein detectable in the AAV9-ABDCon cohort relative to the AAV9 cohort (p=0.0007; Figure 4E). These results show that liver-directed AAV9-ABDCon gene delivery results in more abundant transgene expression from the liver than AAV9.

**Figure 4.**
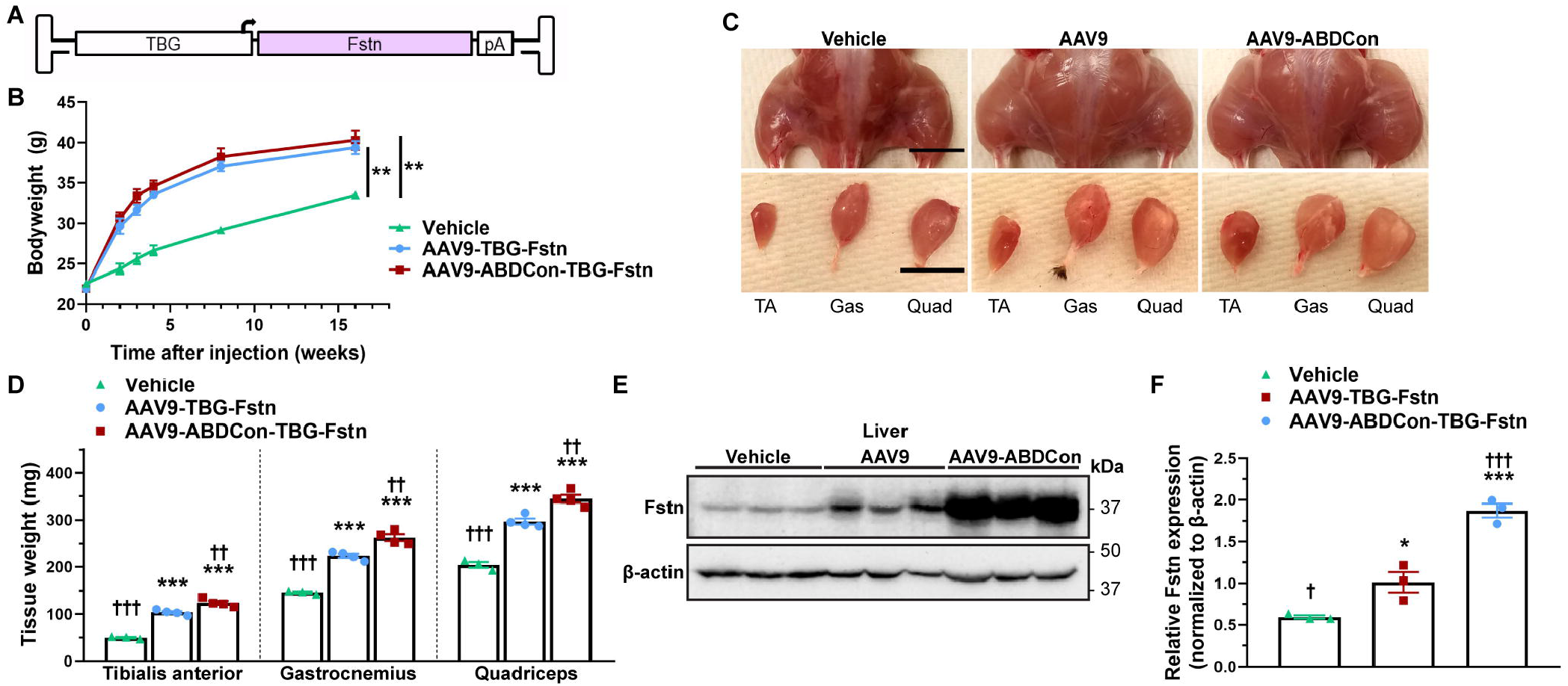
Assessment of AAV9-ABDCon-mediated liver-directed follistatin gene delivery in mice. 6-week-old male C57BL/6J were treated with vehicle (n=3), AAV9-TBG-Fstn (n=4) or AAV9-ABDCon-TBG-Fstn (n=4) at a dose of 5×1012 vg/kg via tail vein injection. **(A)** The TBG-Fstn cassette contains the full-length human codon-optimized Fstn-315 (FS315) sequence under the transcriptional regulation of the liver-specific TBG promoter. **(B)** Bodyweight over time. **(C)** Gross hindlimb musculature and isolated hindlimb muscles. Scale bars represent 1 cm. **(D)** Whole tissue weight of tibialis anterior, gastrocnemius and quadriceps from vehicle- and vector-treated mice. **(E)** Western blot of liver tissue homogenates. The Fstn monomer (38 kDa) was clearly detectable in all samples. β-actin (45 kDa) was used as a loading control. Equal protein loading was verified by Ponceau S staining. **(F)** Western blot densitometry analysis. Fstn expression levels were normalized to β-actin. All error bars represent mean◻±◻SEM. *p◻<◻0.05; **p◻<◻0.01; ***p◻<◻0.001; n.s. not significant; compared to vehicle-treated control. †p◻<◻0.05; †††p◻<◻0.001; compared to AAV9-TBG-Fstn-treated control. TA: tibialis anterior; Gas: gastrocnemius; Quad: quadriceps

## DISCUSSION

Here, we utilize a rational design approach to construct and characterize a modified AAV9 capsid engineered to bind serum albumin. This albumin-binding AAV capsid, called AAV9-ABDCon, is capable of forming intact vector particles without a significant decrease in vector titer. The original purpose of this study was to determine if an albumin-bound AAV9 vector could achieve improved transduction in skeletal muscle in vivo based on the finding by Wang and colleagues that pre-incubation of AAV8 in HSA improves muscle transduction following intramuscular delivery (10). Although we do not observe a noticeable improvement in skeletal muscle transduction, we do see significantly increased liver transduction by AAV9-ABDCon after systemic administration in vivo, a finding which has also been reported in studies evaluating the impact of AAV8 pre-incubation in HSA (10,11). However, in contrast to these prior studies, Denard and colleagues were unable to detect AAV8 binding to HSA, suggesting that a mechanism other than serum albumin-binding may be responsible (21). Altogether, our results suggest serum albumin-binding may be an effective approach to improve the efficiency of AAV vector-mediated gene delivery to the liver, which may be beneficial in the clinical development of improved AAV vectors for liver-directed gene delivery.

The exact mechanism underlying the observed increase in liver transduction by AAV9-ABDCon is unclear, but there are a few potential explanations. The liver is the major site of serum albumin biosynthesis, and albumin-bound complexes have been shown to be directly taken up by hepatocytes (22–24). Additionally, albumin is a well-established ligand of the neonatal Fc receptor (FcRn), and albumin binding to FcRn in the endosome is associated with rescue of albumin from lysosomal degradation. This interaction is thought to explain the unusually long half-life of serum albumin (25). Thus, it may be possible that the observed increased transduction by AAV9-ABDCon is due to increased serum albumin binding-mediated hepatic uptake, FcRn-dependent rescue from lysosomal degradation in the endosome or a combination of these and other potential factors. Additional mechanistic studies are needed to clarify the exact mechanism by which albumin-binding increases AAV-mediated uptake into the liver.

In these studies, the engineered ABDCon peptide, originally developed by Steven Jacobs and colleagues, was selected for its ability to bind serum albumin from several species with high affinity, including human (75 pM), mouse (3.2 nM) and rhesus macaques (61 pM) (12). This cross-species binding may facilitate further preclinical animal study across different animal models. Another advantage of the ABDCon peptide is that serum albumin-binding affinity is tunable and can be easily modulated by specific point mutations in the peptide sequence to generate variants with reduced binding affinity to serum albumin (12). Although not tested here, it is possible that excessively tight binding to serum albumin may impede cellular uptake of the vector, and further studies will be necessary to determine the impact of reduced binding affinity to serum albumin on vector transduction.

It has been demonstrated that a wide variety of functionalized peptides can be inserted at the VP2 N-terminus site to impair additional properties to the AAV capsid, provided the inserted peptide is able to retain its intended conformation (13,14,17,26). The addition of a flexible Gly-Ser linker to attach the peptide insert to the AAV capsid can allow for greater freedom of movement for the conjugated peptide and discourage steric interference from the capsid (27). From our studies, we find that insertion of a 5.4 kDa ABDCon peptide at the AAV9 VP2 N-terminus appears to result in a small but non-significant decrease in titer when produced in adherent HEK293 cells. There is also evidence to suggest that VP2 N-terminal insertions may result in an increase in the empty-to-full capsids ratio (26). These findings indicate that insertions at the AAV VP2 N-terminus may have a mildly detrimental effect on vector production and packaging. As VP2 is nonessential for capsid formation, the final vector preparation would theoretically contain a mixed population of both the desired ABDCon-displaying capsids and capsids containing only VP1 and VP3 (13). This outcome would be expected because it is not possible to ensure every transfected cell receives a copy of both the pVP1,3 and pVP2-ABDCon cap plasmids. Indeed, a large number of purified AAV9-ABDCon capsids were not observed to bind HSA on immuno-TEM. From our studies, it is unclear what fraction of capsids displayed the ABDCon peptide, although Münch and colleagues reported 68.6±10.2% recovery of His-tagged DARPIN-displaying AAV capsids using a HisTrap affinity chromatography column (28). We predict a similar chromatographic approach using HSA as the affinity ligand could be used to isolate and purify AAV9-ABDCon capsids.

Although we observe a robust increase in murine liver transduction following intravenous delivery with the AAV9-ABDCon vector, it is unclear if this would translate to a similar effect in non-human primates or humans. It has become evident that AAV transduction profiles across different species and, in exceptional cases, different strains may vary drastically (6,29). Thus, a prudent next step would be evaluation of AAV9-ABDCon in different mouse strains along with assessment in large animal models. In addition, it would be worthwhile to assess the feasibility of serum albumin-binding peptide insertion on other AAV capsids, such as the human hepatocyte-tropic AAV-LK03 (6).

## ACKNOWLEDGEMENTS

We thank Aravind Asokan for providing the capsid B1 antibody and Victoria Madden for assistance in planning, executing and interpreting immuno-TEM experiments.

## AUTHOR CONTRIBUTIONS

QJ and XX conceived and planned the experiments. QJ, CQ and Jianbin Li carried out the experiments. QJ performed analysis of the data. QJ, Juan Li and XX contributed to the interpretation of the results. QJ prepared the figures and drafted the manuscript. XX procured funding support. All authors reviewed and provided critical feedback on the final manuscript.

## CONFLICT OF INTEREST

XX and QJ are co-inventors on a potential patent application related to the work. The remaining authors declare that they have no conflict of interest.

## FUNDING

This work was funded by an award from the Eshelman Institute for Innovation (EII) to XX.

