## Supplemental Methods for "An engineered serum albumin-binding AAV9 capsid achieves improved liver transduction after intravenous delivery in mice"

**Supplementary Methods**

**Generation of modified AAV9-ABDCon rep/cap plasmids**. The mutated AAV9 cap gene encoding unmodified VP1/VP3 (pVP1/3) was generated by removing the VP2 start site on the AAV9 rep/cap plasmid by site-directed mutagenesis (T138A). T138A primers: forward 5' GCCGCTCCTGGAAAGAAGAG 3' and reverse 5' CTTAGCCGCTTCCTCAACCAG 3'. For generation of the modified cap gene containing the VP2 N-terminal ABDCon peptide insertion, the codon-optimized DNA sequence encoding for the ABDCon peptide preceded by a Kozak consensus sequence, GCCACCATG, was inserted at the N-terminus of the AAV9 VP2 on the AAV9 cap gene. The entire modified region including the HindIII/ApaI restriction site on the AAV9 cap gene were synthesized (GenScript; Piscataway, NJ) and cloned into the AAV2/9 rep/cap plasmid. To prevent erroneous translation of a modified VP1 containing the ABDCon peptide, the VP1 start site was mutated to leucine (M1L). M1L primers: forward 5' CTGGCTGCCGATGGTTATCTTC 3' and reverse 5' ACCTGGTTTAAGTCATTTATTGTTCAG 3'.

**AAV vector production**. AAV9 and AAV9-ABDCon vectors were produced by triple plasmid transfection. To supply AAV9-ABDCon rep/cap sequences, pVP1/3 and pVP2-ABDCon were transfected in a 5:1 final molar ratio. 48-72 h after transfection, AAV vectors were purified from HEK293 lysates and media by polyethylene glycol (PEG) precipitation and two rounds of cesium chloride (CsCl) density ultracentrifugation. Fractions containing AAV were dialyzed overnight to remove residual CsCl and purified vectors were stored at -80°C.

**Quantitative DNA dot blot**. Briefly, purified AAV vectors underwent DNAse treatment to remove unencapsulated DNA followed by proteinase K treatment to open capsid particles. Vector DNA was precipitated using 100% ethanol and washed with 70% ethanol. The DNA pellet was dissolved in alkaline buffer (0.4M NaOH, 10 mM EDTA, pH 8.0) and blotted on a positively-charged nylon membrane. To detect vector DNA, the membrane was incubated at 55°C with a biotinylated probe designed to hybridize to sequences on the packaged transgene cassette. Streptavidin-HRP was added to the membrane at room temperature and signals were developed with North2South Stable Peroxide Solution and Luminol/Enhancer Solution (Thermo Scientific; Rockford, IL). Densitometry analysis was conducted in AlphaView (ProteinSimple; San Jose, CA). Biotinylated probes were generated using the North2South Biotin Random Prime Labeling Kit (Thermo Scientific; Rockford, IL). Vector titers were quantified at least twice using at least two different biotinylated probes corresponding to distinct regions (typically the promoter and the transgene sequence) on the transgene cassette.

**SDS-PAGE and western blot.** For western blot of AAV capsid proteins, purified AAV vectors were boiled in Laemmli sample buffer (62.5 mM Tris-HCl, 1.5% SDS, 8.3% glycerol, 1.5% β-mercaptoethanol, 0.005% bromophenol blue, pH 6.8) and run on a 10% SDS-PAGE gel. A mouse monoclonal anti-AAV VP1/VP2/VP3 (1:1000; B1 antibody) was used for detection of capsid proteins. For western blot of liver tissue, 30 µg of liver tissue homogenate was loaded on a 10% SDS-PAGE gel. Equal protein loading was verified by Ponceau S staining. The antibodies used were rabbit polyclonal anti-follistatin (1:1000; H-114, sc-30194; Santa Cruz Biotechnology; Dallas, TX) and HRP-conjugated mouse monoclonal anti-β-actin (1:25,000; AC-15, A3854; Sigma Aldrich; St. Louis, MO). Western blot densitometry analysis was conducted in AlphaView software (ProteinSimple; San Jose, CA).

**Alkaline agarose gel electrophoresis.** Briefly, purified AAV vectors were treated with DNAse followed by proteinase K. Vector DNA was separated by phenol/chloroform extraction and precipitated with ethanol. The DNA pellet was resuspended in TE (10 mM Tris-HCl, 2 mM EDTA, pH 8.0) and samples were run on a 1% alkaline agarose gel (50 mM NaOH, 2 mM EDTA) in alkaline agarose gel running buffer (50 mM NaOH, 2 mM EDTA). After the run, the gel was neutralized with 1X TAE and stained in ethidium bromide in 1X TAE overnight at 4°C.

**Vector genome quantification.** Total DNA was extracted from tissue samples using DNeasy Blood & Tissue Kit (Qiagen; Hilden, Germany) according to the manufacturer's instructions. The number of vector copies was determined by absolute qPCR quantification using primers and a TaqMan probe sequence recognizing the vector-derived CB promoter sequence. The number of vector copies was normalized to the endogenous mouse glucagon gene. Assays were performed using GoTaq PCR Master Mix (Promega; Madison, WI) on a 7300 Real-Time PCR System (Applied Biosystems; Foster City, CA). Primers and probe sequences: CB forward 5' GTATGTTCCCATAGTAACGCCAATAG 3'; CB reverse 5' GGCGTACTTGGCATATGATACACT 3'; CB probe 5' FAM-TCAATGGGTGGAGTATTTA-MGB 3'; glucagon forward; 5' AAGGGACCTTTACCAGTGATGTG 3'; glucagon reverse 5' ACTTACTCTCGCCTTCCTCGG 3'; glucagon probe 5' FAM-CAGCAAAGGAATTCA-MGB 3'
